# A Microsporidian blocks *Plasmodium falciparum* transmission in *Anopheles arabiensis* mosquitoes

**DOI:** 10.1101/799445

**Authors:** Jeremy K. Herren, Lilian Mbaisi, Enock Mararo, Joseph W. Oundo, Edward E. Makhulu, Hellen Butungi, Maria Vittoria Mancini, Victor A. Mobegi, Jordan Jabara, Steven P. Sinkins

## Abstract

Malaria imposes an enormous burden on sub-Saharan Africa, and evidence that incidence could be starting to increase again^1^ suggests the limits of currently applied control strategies have now been reached. A possible novel control approach involves the dissemination in mosquitoes of inherited symbiotic microbes to block transmission. This strategy is exemplified by the use of transmission-blocking *Wolbachia* in *Aedes aegypti* against dengue virus^2–7^. However, in the *Anopheles gambiae* complex, the primary African vectors of malaria, there limited reports of inherited symbionts with transmission-blocking capacity^8–10^. Here we show that a newly discovered vertically transmitted species of *Microsporidia* symbiont in the *An. gambiae* complex blocks *Plasmodium* transmission. *Microsporidia MB* is present at moderate prevalence in geographically dispersed populations of *An. arabienesis* in Kenya, localized to the mosquito midgut and ovaries, and is not associated with significant reductions in adult host fecundity or survival. Field collected *Microsporidia MB*-infected *An. arabiensis* were never found to harbor *P. falciparum* gametocytes and on experimental infection with *P. falciparum* no sporozoites could be detected in *Microsporidia MB-*infected mosquitos. As a *Plasmodium* transmission-blocking microbe that is non-virulent and vertically transmitted, *Microsporidia MB* could be exploited as a novel malaria control tool.

---

Microsporidia are a group of obligately intracellular simple eukaryotes, classified within or as a sister group to fungi, and found in a wide range of hosts, but most commonly invertebrates. Their lifecycles include a meront phase during proliferation, and spores with chitinous cell walls involved in host to host transmission through spore ingestion. Species with solely horizontal transmission usually show greater virulence and lower host specificity, but where a combination of horizontal and vertical transovarial transmission occurs, lower virulence is advantageous, and is normally associated with a higher degree of host specificity^11–12^. Sex ratio distortion toward females has been reported (a manipulation characteristic of transovarially transmitted symbionts), for example *Dictyocoela* microsporidia in Amphipod crustaceans^13^. Various microsporidia species have been reported in mosquitoes^14–25^, with simple or complex lifecycles^14^ but all of which are pathogens where virulence is primarily associated with larval mortality or reduced adult fecundity and lifespan^14–20^.

In this study a previously unknown species of *Microsporidia* was discovered during microbiome characterization of *An. arabiensis*, designated *Microsporidia MB*, and was found to occur at a high density and low to moderate prevalence (0-10%) in geographically dispersed populations of *An. arabiensis* in Kenya. Notably, in all *An. arabiensis* populations investigated, we observed that none of the *Microsporidia MB* harboring mosquitoes were infected with *Plasmodium* (Extended Data Fig. 1). Phylogenetic analysis of the 18S ribosomal gene revealed that *Microsporidia MB* is related to *Crispospora chironomi*^26^, a species recently identified from non-biting midges. The 18S gene sequence of *Microsporidia MB* shows 97% similarity with *Crispospora chironomi*. *Microsporidia MB* and *Crispospora chironomi* are in clade IV that unites microsporidia of terrestrial origin infecting diverse hosts^27^ (Fig. 1A and Extended data Fig. 1). The previous reports of microsporidia infecting *Anopheles* mosquitoes all belong to different clades of *Microsporidia*. The morphology of *Microsporidia MB* closely resembles *Crispospora chironomi*, exhibiting both polysporoblastic and diplosporoblastic sporogenies, both found in the larval mosquito gut epithelium (Fig. 1B).

**Figure 1:**
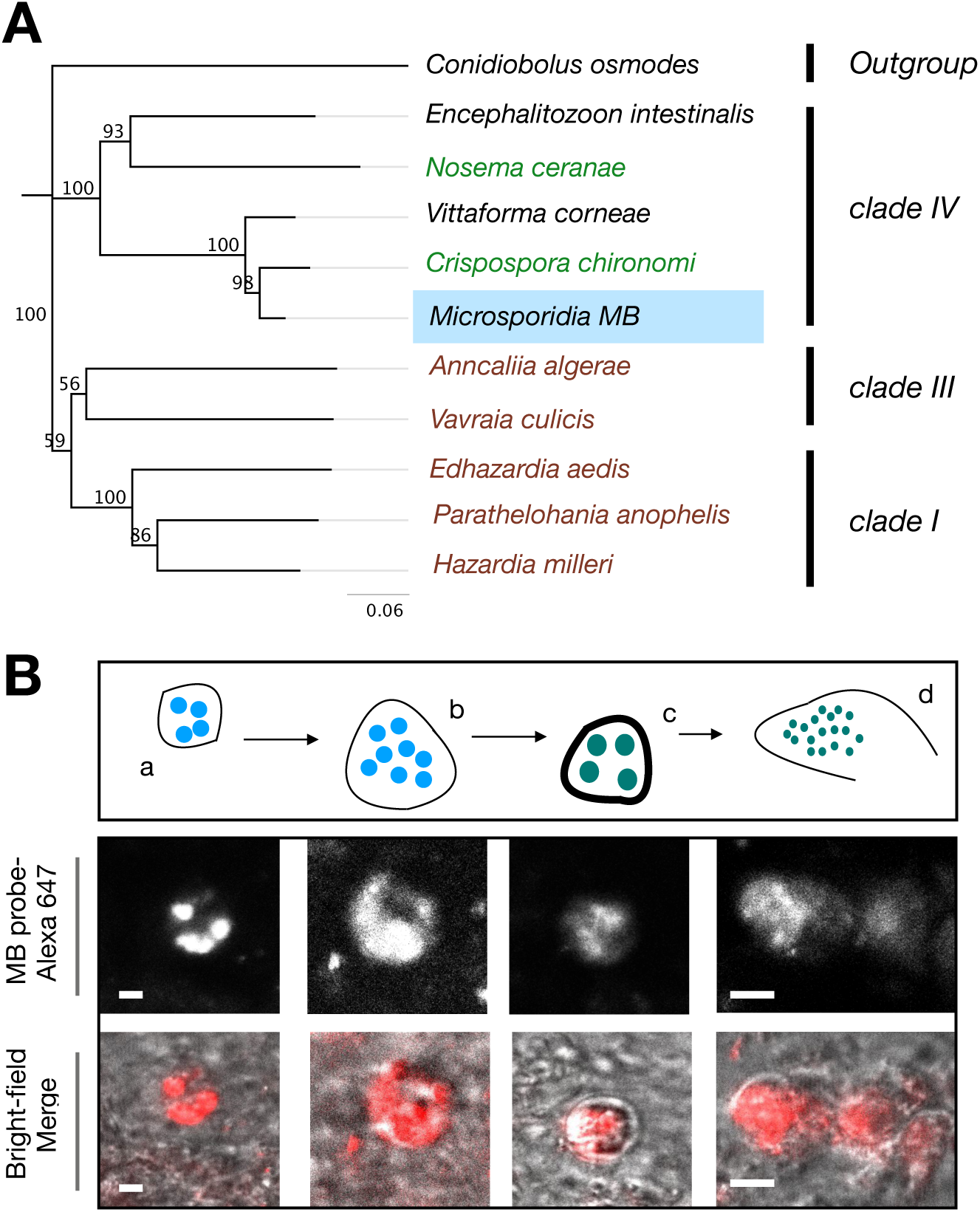
Novel Microsporidian associated with *An. arabiensis* populations in Kenya. (A) 18S rDNA-based phylogeny reveals that *Microsporidia MB* are in clade IV of the Microsporidia. Labelled in brown are the microsporidian species known from mosquitoes. In green are microsporidians associated with other insect groups. (B) FISH staining of the diplokaryotic stages of *Microsporidia MB* merogony (a-b) and spore capsule formation (c-d) in *An. arabiensis* larval gut epithelial tissues. Scale bar,1.5 μm.

The effects of *Microsporidia MB* on *Plasmodium* infection in *An. arabiensis* was further examined around Mbita point, western Kenya, using G_1_ offspring pools obtained from field-collected G_0_ mosquitoes and membrane feeding assays (MFAs) carried out with *Plasmodium falciparum* infected donor blood. Since there was a low to moderate prevalence of *Microsporidia MB* is field populations (Extended Data Fig. 2), we screened mosquitoes for *Microsporidia MB* and sorted them to ensure that G_1_ progeny pools used for MFAs would have both *Microsporidia MB*-infected and uninfected mosquitoes. *Plasmodium* was quantified 10 days after MFAs, and a strong negative correlation was apparent between the *Microsporidia MB* and the presence of *Plasmodium* in whole mosquitoes 10 days after MFA (Fig. 2A-B). *Plasmodium* parasites in an ingested bloodmeal undergo a series of developmental changes prior to traversing the peritrophic matrix and midgut epithelium to form a sporogenic oocyst, which releases sporozoites into the hemocoel. From the hemocoel, sporozoites travel to the salivary gland, traverse an epithelium and mix with *Anopheles* saliva resulting in an infectious mosquito, usually 8-14 days after the blood meal^28^. To investigate the stage at which *Plasmodium* development was inhibited, we specifically quantified *Plasmodium* in the *Anopheles arabiensis* head and thorax and abdominal compartments. The absence of *Plasmodium* in the head and thorax compartment of *Microsporidia MB* infected *Anopheles arabiensis* 10 days after MFA indicates that *Microsporidia MB* prevents *Anopheles arabiensis* salivary glands from being colonized by *Plasmodium* sporozoites, and therefore prevents *Plasmodium* transmission (Fig. 2C-D). In addition, no *Plasmodium* oocysts are established in the midgut of *Microsporidia MB* infected *Anopheles arabiensis* as determined by the absence of *Plasmodium* infections in abdomens 10 days after MFA (Extended data Fig. 3). These results indicate that *Microsporidia MB*-induced transmission blocking occurs prior to the establishment of oocysts in the *Anopheles* mosquito midgut.

**Figure 2:**
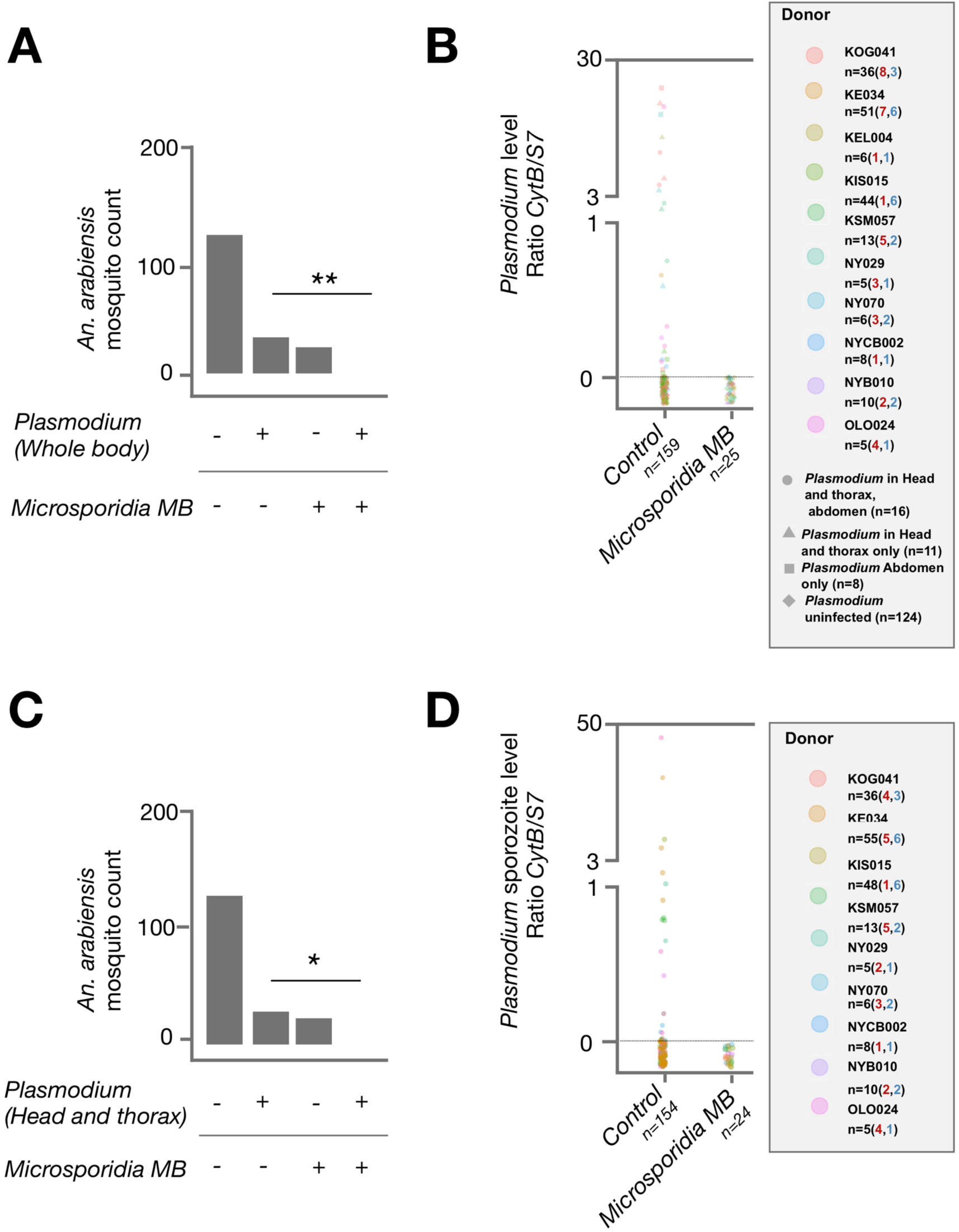
*Microsporidia MB* blocks parasite development in *An. arabiensis* after membrane feeding assay challenge with *P. falciparum*. (A) The *Plasmodium* infection rate in *Microsporidia MB* positive and *Microsporidia MB* negative mosquitoes was determined by qPCR. There was a significant absence of co-infected mosquitoes (two tailed fisher exact test, P=0.005 N=184). (B) *Plasmodium* density quantified by qPCR in *An. arabiensis*. For *An. arabiensis* mosquitos infected in both head and thorax and abdomen, average *Plasmodium* density across both compartments is given. (C) The head and thorax *Plasmodium* infection rate, reflecting presence of sporozoites, in *Microsporidia MB* positive and *Microsporidia MB* negative mosquitoes. There was a significant absence of co-infected mosquitoes (two tailed fisher exact test, P=0.02 N=178). (D) *Plasmodium* density in *An. arabiensis* heads and thoraxes, quantified by qPCR in. Data shown in A,B, C, D is pooled from replicate experiments carried out using different gametocyte donors (for each donor the numbers of *Plasmodium* positive *An. arabiensis* is shown in red and *Microsporidia MB* positives *An. arabiensis* is shown in blue). Each data point is an individual *An. arabiensis* mosquito. If *Plasmodium* was detected in either head and thorax or abdomens or both compartments, the *An. arabiensis* was considered to have a *Plasmodium* infection (A,B). Heads and thoraxes were separated and screened individually for head and thorax specific infection rate (C,D).

**Figure 3:**
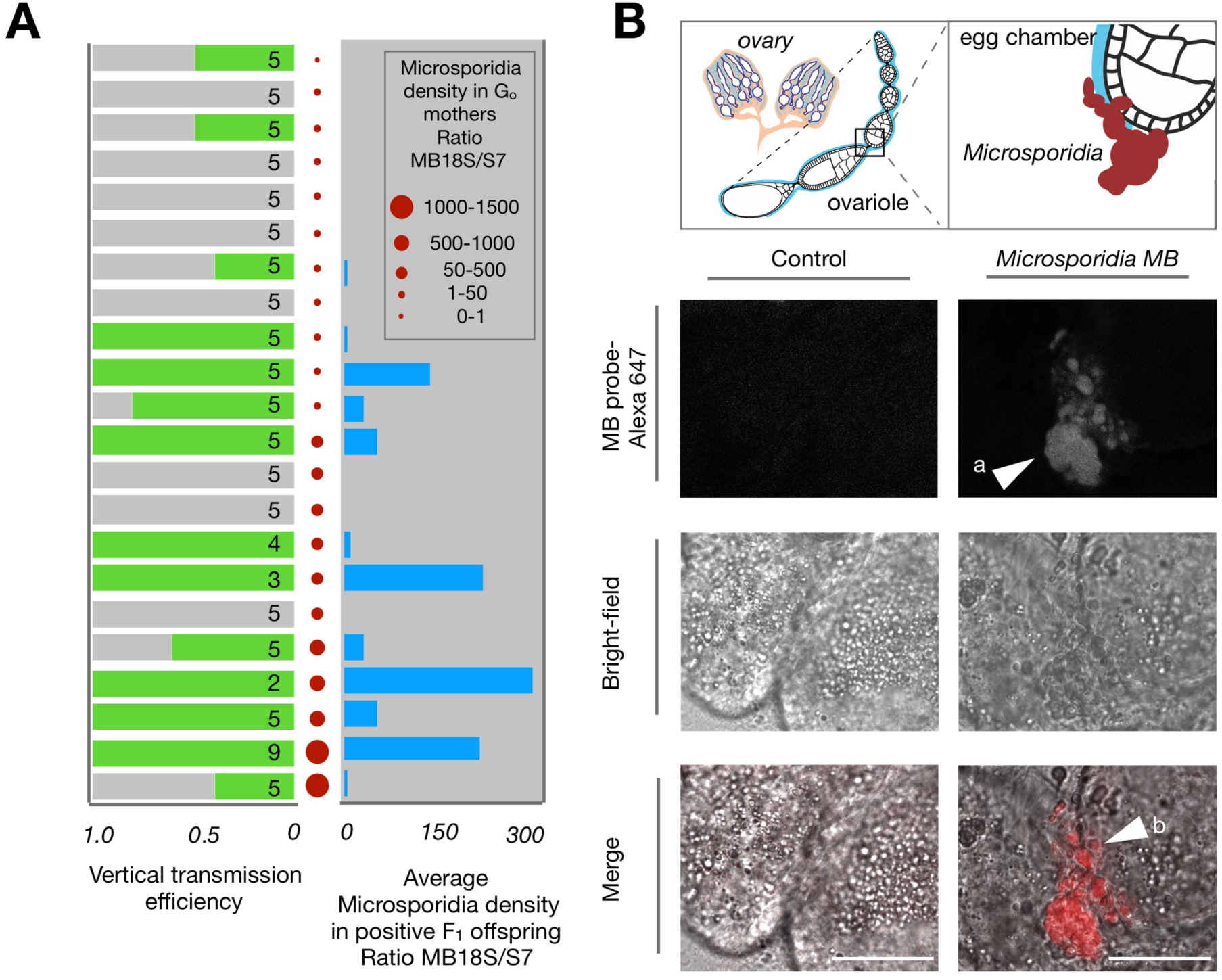
*Microsporidia MB* is maternally transmitted. (A) The vertical transmission efficiency and density of *Microsporidia*^*MB*^ in G_1_s depends on the G_0_ (maternal) *Microsporidia MB* density. Each red dot represents a *Microsporidia MB*-infected G_0_ female *Anopheles arabiensis* mosquito. The density of *Microsporidia MB* in G_0_ females is reflected by the size of the red dot. Green bars depict the vertical transmission efficiency from G_0_ to G_1_, whereas blue bars depict the density of *Microsporidia MB* in G_1_s. Numbers of G_1_s tested to determine vertical transmission efficiency and *Microsporidia MB* density are given in the bars. (B) Fluorescence microscopy with a *Microsporidia MB*-specific FISH probe (MB probe) indicates that *Microsporidia MB* (a) is localized to the posterior of developing vitellogenic egg chambers (b) in *An. arabiensis*. Scale bar, 20 μm.

*Microsporidia MB* are generally maternally transmitted with high efficiency (Fig. 3A), from 45-100%. However, a number of wild-caught females with low *Microsporidia MB* density did not transmit or transmitted very poorly to their offspring. It is possible that these infections might be newly acquired and have not yet become localized to the ovaries, a likely requirement for high-efficiency maternal transmission. *Microsporidia MB* were observed in the mosquito ovaries, where the symbiont colonizes and penetrates oocytes (Fig. 3B).

The fecundity and egg to adult survival rate of *Microsporidia MB* infected and uninfected iso-female lineages were then examined. No significant differences were observed in the number of eggs laid by *Microsporidia MB* infected versus uninfected individuals, indicating that *Microsporidia MB* does not have a sterilizing effect on females (Fig. 4A). A shortened development period from egg to adult was observed for individuals carrying *Microsporidia MB* (Fig. 4B). We investigated the survival of adult female mosquitoes harboring *Microsporidia MB* and found their longevity was similar to uninfected mosquitoes (Extended Data Fig. 3). The density of *Microsporidia MB* was examined across the lifecycle of the mosquito and was higher in adults than in larvae, but larval density is highly variable. Notably, *Microsporidia MB* levels are lower in recently emerged adults, increasing with mosquito age (Extended Data Fig. 3).

**Figure 4:**
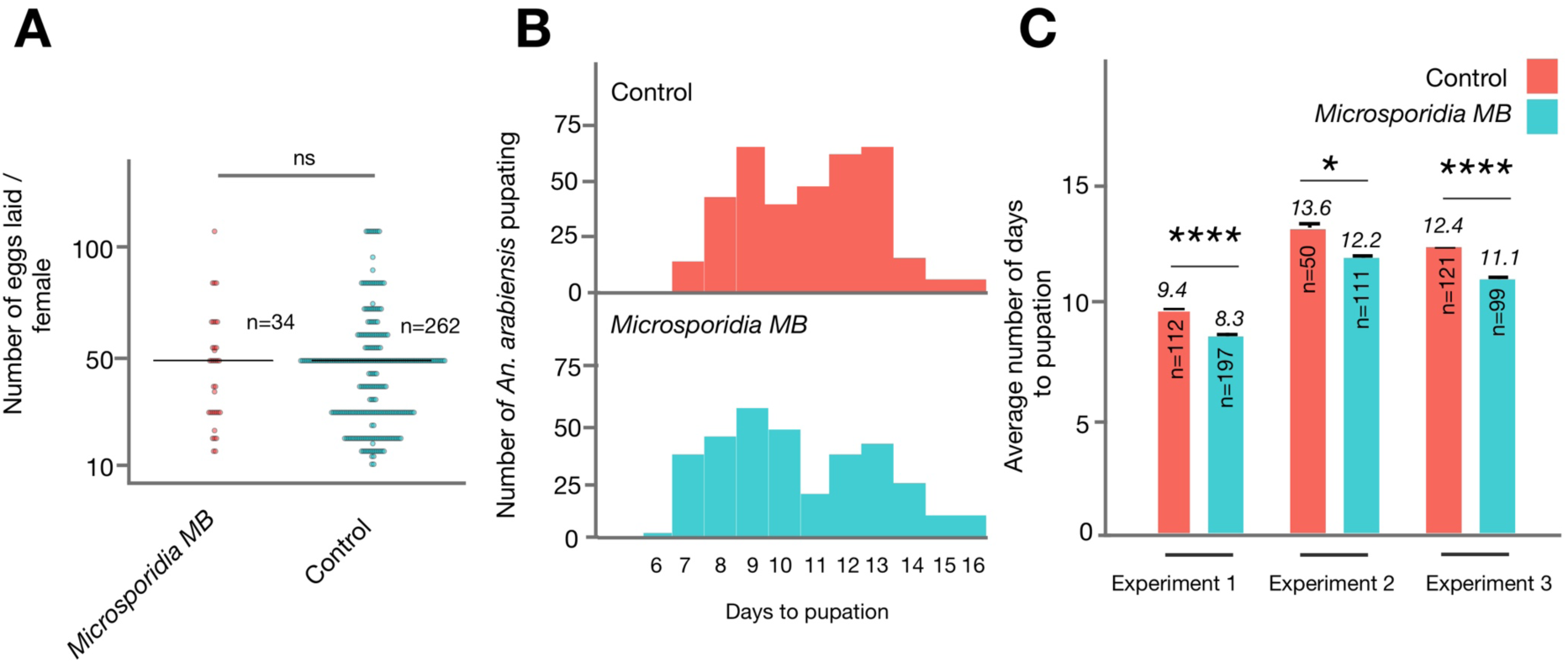
*Microsporidia MB* does not overtly decrease host fitness. (A) *Microsporidia MB*-harboring wild-caught mosquitoes did not lay significantly less eggs than uninfected counterparts. Black line indicates means. (B) The F_1_ larval progeny of *Microsporidia MB*-infected wild-caught females develop significantly faster than uninfected counterparts, data shown is pooled from three independent experiments. (C) The larval development of *Microsporidia MB*-infected *An. arabiensis* is on average 1.1, 1.4 and 1.3 days less than uninfected controls in three independent experiments, error bars reflect SEM.

Several *Anopheles*-associated microsporidians have been shown to interfere with the infection and development of *Plasmodium*. *Nosema stegomyiae* disrupts the development of the oocysts in *An. gambiae*, attributed to mid-gut degradation and consequent disruption of *Plasmodium* binding^24^, while *Vavraia culicis* inhibition of development of *Plasmodium* has been associated with host innate immune priming^25^. The inhibitory effects observed in previous *Plasmodium* transmission experiments have been relatively modest, with only partial reduction of transmission observed. In applied terms, given their virulence, their primary application would be as population suppression agents and the *Plasmodium* inhibition would provide a minor add-on. In contrast, complete *P. falciparum* transmission blocking was observed here for *Microsporidia MB*. Additionally, *Microsporidia MB* blocking occurs early, prior to the formation of *Plasmodium* oocysts in the *Anopheline* mosquito gut. When combined with the key characteristics that will facilitate artificially elevating its population frequency, namely spore production that is likely to facilitate dissemination, efficient transovarial transmission and apparently non-virulent interactions with *An. arabiensis* mosquitoes, *Microsporidia MB* is a realistic candidate for a stable vector population replacement strategy for malaria control.

These findings are significant in terms of regional malaria transmission and epidemiology as well as risk-mapping, and particularly in terms of the development of microbe-based *Plasmodium* transmission blocking tools. *Microsporidia MB* can be added to the *Anopheles*-associated gut bacteria that have previously been identified as able to reduce *Plasmodium* transmission when introduced into the mosquito^29,30^, or following introduction of anti-*Plasmodium* transgenes^31^. As an unmodified, *Anopheles*-associated inherited endosymbiont that confers highly effective malaria blocking under natural conditions, it is a promising prospect for malaria control.

## Materials and Methods

### Ethics statement

Ethical clearance (Kenya Medical Research Institute Scientific and Ethics Review Unit: KEMRI/RES/7/3/1 and Glasgow MLVS College Ethics Committee: Project Number 200170001) was obtained prior to human blood sample collection. Written informed consent was sought from parents and guardians of the children to allow minors to participate in the study. Consent was also obtained from heads of households that provided approval for indoor mosquito collection.

### Data and materials availability statement

All data is available in the main text or the extended data.

### Sampling sites and collection

*Microsporidia MB* and *Plasmodium* prevalence in wild *Anopheles arabiensis* mosquitoes was determined by collecting adult female mosquitoes from sites around Kenya: Mbita (Nyawiya, Mageta and Kirindo), Mwea (Mbui-Njeru), Busia (Funyala) and Ahero (Kigoche). *An. arabiensis* were collected inside houses and sheds using CDC light traps and by manual aspiration. For the establishment of *Microsporidia MB* harboring lines gravid mosquitoes were collected solely by manual aspiration inside houses and sheds. All mosquitoes were transported from the field to the *icipe*-TOC laboratories and insectaries alive in cages, each sample represents an individual wild caught mosquito.

### Mosquito species identification

All experiments were carried out on wild collected *Anopheles gambiae sl.*, which were identified morphologically. In all of the collection sites, *Anopheles arabiensis* is the most common member of the *An. gambiae* species complex, with >97% of complex members being identified as *Anopheles arabiensis*. The species designation was confirmed using a molecular assay that differentiates *An. gambiae s.s.* and *An. arabiensis* using the SINE S200 X6.1 locus^31^. *Anopheles* samples that were not confirmed to be *Anopheles arabiensis* were excluded from analysis.

### Determination of Molecular Phylogeny of *Microsporidia MB*

*Microsporidia MB* positive *An. arabiensis* were initially identified by sequencing 18S amplicons amplified by the SSU rRNA primer pair F: 5′-CACCAGGTTGATTCTGCC-3′; R: 5′-TTATGATCCTGCTAATGGTTC-3′, which targets phylogenetically diverse microsporidians^32^. The primer pair RPOBMBF 5′-ACAGTAGGTCACTTGATTGAATGTC-3′ and RPOBMBR 5′-TACCATGTGCTTAAGTCTTTGGT-3′ was used to amplify the *rpoB* gene of *Microsporidia MB*. Amplicaons were prepared from nine individual *An. arabiensis* from geographically dispersed sites, all individuals had identical 18S and *rpoB* fragment gene sequences. Prepared amplicons were cleaned using the USB^®^ ExoSAP-IT^®^ PCR Product Cleanup kit according to manufacturer’s instructions and sent to Macrogen (Netherlands) for sequencing. Multiple sequence alignment was done using the MUSCLE algorithm^33^ alongside reference sequences of other *Microsporidia* species obtained from NCBI. Tamura-Nei genetic distance model alongside Neighbour-joining tree building algorithm was used in the creation of phylogenies and evaluated with 10000 replicates bootstrap support and 50% support threshold. Rooting was done using *Conidiobolus osmodes* as an outgroup. *Microsporidia MB* 18S rDNA and *rpoB* partial gene reference sequences have been submitted to Genbank, submission ID 2270798.

### Egg laying and establishment of lines

Wild-caught gravid female mosquitoes were induced to oviposit inside a perforated 1.5ml micro centrifuge tube containing 50µl of distilled water and a soaked piece of Whatman paper towel. Eggs from each female were counted under a compound microscope using a paint brush and then dispensed into water tubs for larval development under optimal rearing conditions (a temperature of 30.5 °C and 30% humidity). Upon laying eggs, the G_0_ females were screened for presence of *Microsporidia MB* by PCR. The number of eggs laid by *Microsporidia MB* infected and uninfected mosquitoes was recorded and two-tailed Mann Whitney U tests were used to determine statistical significance. The progeny from *Microsporidia MB* positive G_0_ females were used for: i) the quantification of vertical transmission efficiency and ii) the establishment of iso-female *Microsporidia MB* infected lines, whereas lines with similar L1 larvae numbers (+/− 10 individuals) were used as matched negative the controls for quantifying development time and survival. For each iso-female line two G_1_ L4 larvae were screened to determine species and confirm infection status and the line was only considered infected if both were *Microsporidia MB* positive. For the quantification of density across developmental stages five infected iso-female lines were established, two of which were monitored until day eight and three of which were monitored until day fifteen. All the mosquitoes in each experiment were derived from a single infected female. To establish mixed (*Microsporidia MB* positive and negative) *An. arabiensis* pools for standard membrane feeding assays, eggs from *Microsporidia MB-*infected and uninfected females were combined and reared together at an approximate ratio of 3:1 (*Microsporidia MB infected*: *Microsporidia MB uninfected*). Infection status was not determined at the larval stage for mixed pools.

### Mosquito rearing

Larvae were reared under a controlled environment at 30.5 °C (+/− 2°C). Larvae were fed daily on TetraMin™ baby fish food and fresh double-distilled water added into their tubs every other day to maintain oxygen levels. Adult mosquitoes were reared at 30°C (+/− 2°C) and 70% humidity with a constant 12-hour day / night cycle. The adult mosquitoes were fed on 6% glucose soaked in cotton wool. For adult survival and larval development time measurements, the status of larvae and mosquitoes was recorded every 24hrs, the Log-Rank and two-tailed Mann Whitney U tests were used to determine levels of statistical significance, respectively.

### *Plasmodium* standard membrane feeding assays

*Plasmodium* screening of human subjects was done in the regions surrounding Mbita using RDT kits (SD Bioline, UK). Microscopy was carried out on RDT-positive samples to confirm the presence of *P. falciparum* gametocytes. Gametocyte-positive blood used was mixed with an anticoagulant (heparin) and a total volume of 100µl was placed into mosquito mini-feeders at 37°C and covered in stretched parafilm. Mixed pools of 2-3 day old *An. arabiensis* (containing co-reared *Microsporidia MB* positive and negative mosquitoes) were starved for 5 hours (the sucrose source was replaced with water for the first 4 hours of starvation) prior to the standard membrane feeding assay. Mosquitoes were allowed to feed for an hour after which non blood-fed individuals were discarded. Blood-fed mosquitoes were then maintained for a period of 10 days post-infection and processed for the molecular detection of *Microsporidia MB* and *Plasmodium* oocysts and sporozoites. We only include experiments where there was greater than zero prevalence of *Microsporidia MB* and *Plasmodium* in the mixed pools of *Anopheles arabiensis.* Two-tailed fisher exact tests were used to determine statistical significance. Each data point represents an individual mosquito.

### Specimen storage and DNA extraction

All *An. arabiensis* specimens were dry frozen at −20°C in individual microcentrifuge tubes prior to DNA extraction. Prior to extraction, mosquitoes were sectioned into head and thorax (for detection and quantification of *Plasmodium* sporozoites) and abdomens (for detection and quantification of *Plasmodium* oocysts and *Microsporidia MB*). DNA was extracted from each section individually using the protein precipitation method (Puregene, Qiagen, Netherlands).

### Molecular detection and quantification of *Microsporidia MB*

*Microsporidia MB*-specific primers (MB18SF: CGCCGGCCGTGAAAAATTTA and MB18SR: CCTTGGACGTGGGAGCTATC) were designed to target the *Microsporidia MB* 18S rRNA gene region and tested for specificity on a variety of Microsporidia-infected mosquito controls (including *Hazardia, Parathelohania* and *Takaokaspora*). For detection, the PCR reaction volume was 10µl, consisting of 2µl HOTFirepol® Blend Master mix Ready-To-Load (Solis Biodyne, Estonia, mix composition: 7.5 mM Magnesium chloride, 2mM of each dNTPs, HOT FIREPol® DNA polymerase), 0.5µl of 5 pmol/µl of both forward and reverse primers, 2 µl of the template and 5 µl nuclease-free PCR water. The PCR cyclic conditions used were; initial denaturation at 95°C for 15 minutes, further denaturation at 95°C for 1 minute, followed by annealing at 62°C for 90 seconds and extension at 72°C for a further 60 seconds, all done for 35 cycles. Final elongation was done at 72°C for 5 minutes. *Microsporidia MB* was also quantified by qPCR using MB18SF/ MB18SR primers, with normalization against the *Anopheles* ribosomal S7 host gene (primers, S7UF: GMCGGGTCTGWACCTTCTGG and S7UR: TCCTGGAGCTGGAARTGAAC). The qPCR reaction volume was 10µl, consisting of 2 µl HOT FIREPol® EvaGreen® HRM no ROX Mix (Solis Biodyne, Estonia, mix composition: 12.5 mM Magnesium chloride, EvaGreen® dye, BSA, dNTPs, HOT FIREPol® DNA Polymerase and 5× EvaGreen® HRM buffer), 0.5µl of 5 pmol/µl of both forward and reverse primers, 2 µl of the template and 5 µl nuclease-free PCR water. Finally, a melt curves were generated including temperature ranges from 65°C to 95°C. Standard curves were also generated to determine amplification efficiency.

### Molecular detection and quantification of *Plasmodium*

A qPCR-based assay was used to detect and quantify the *cytochrome b* gene of *Plasmodium*^34^. When utilized on *An. arabiensis* head and thorax DNA samples 10 days post MFA this assay can be used to detect and quantify *Plasmodium* sporozoites. When utilized on *An. arabiensis* abdomen DNA samples 10 days post MFA this assay can be used to detect and quantify *Plasmodium* oocysts. Plasmodium *cytochrome b* was normalized against the *Anopheles* ribosomal S7 host gene. The qPCR mastermix was composed of 2 µl HOT FIREPol^®^ EvaGreen^®^ HRM no ROX Mix (Solis Biodyne, Estonia, mix composition: 12.5 mM Magnesium chloride, EvaGreen^®^ dye, BSA, dNTPs, HOT FIREPol^®^ DNA Polymerase and 5× EvaGreen^®^ HRM buffer), 0.5 µl of 5 pmol/µl of both forward (cytBF) and reverse (cytBR) primers, 2 µl of the template and 5 µl nuclease-free PCR water. The PCR profile for target gene amplification included an initial denaturation at 95 °C for 15 minutes, further denaturation at 95 °C for 30 seconds, followed by annealing at 60 °C for 45 seconds and extension at 72 °C for 45 seconds repeated for 40 cycles. Final elongation was performed at 72 °C for 7 minutes followed by generation of a melt curve ranging from 65 °C to 95 °C. A standard curve was generated to determine the PCR efficiency.

### Fluorescence *in situ* Hybridisation (FISH)

DNA probes specific to *Microsporidia MB* 18S rDNA were designed and synthesized (5’-CY5-CCCTGTCCACTATACCTAATGAACAT-3’, Macrogen, Netherlands). FISH was conducted on *An. arabiensis* adult and larval tissue specimens using a previously described protocol^35^, with minor modifications. Briefly, mosquito tissues (larval gut and adult ovaries) were fixed in 4% Paraformaldehyde (PFA) solution overnight at 4°C and subsequently transferred into 10% hydrogen peroxide in 6% alcohol for 3 days prior to rehydration in Phosphate Buffer Saline with Tween-20 (PBS-T) for 1–2 hours. Hybridization was conducted by incubating tissues in 150 μl hybridization buffer (20 mM Tris-HCl [pH 8.0], 0.9M NaCl, 0.01% sodium dodecyl sulfate, 30% formamide) containing 100 pmol/ml of the probe at room temperature overnight. After washing with PBS-T, the hybridized samples were placed on a slide and were visualized immediately using a Leica SP5 confocal microscope (Leica Microsystems, USA).

## Extended Data

**Extended Data Figure 1:**
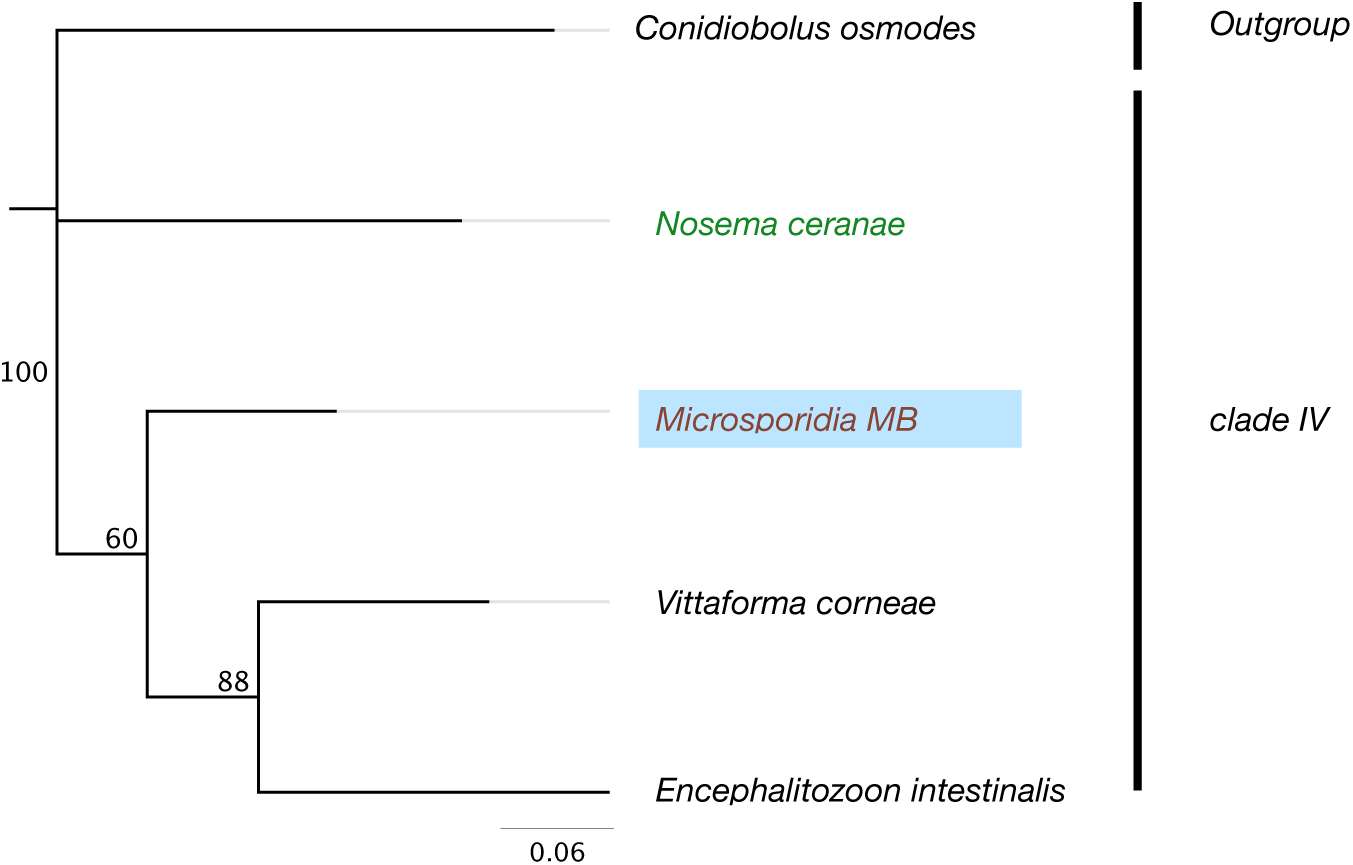
The RNA polymerase B’ subunit gene (*rpoB*) based phylogeny reveals that Microsporidia MB are in clade IV of the Microsporidia.

**Extended Data Figure 2:**
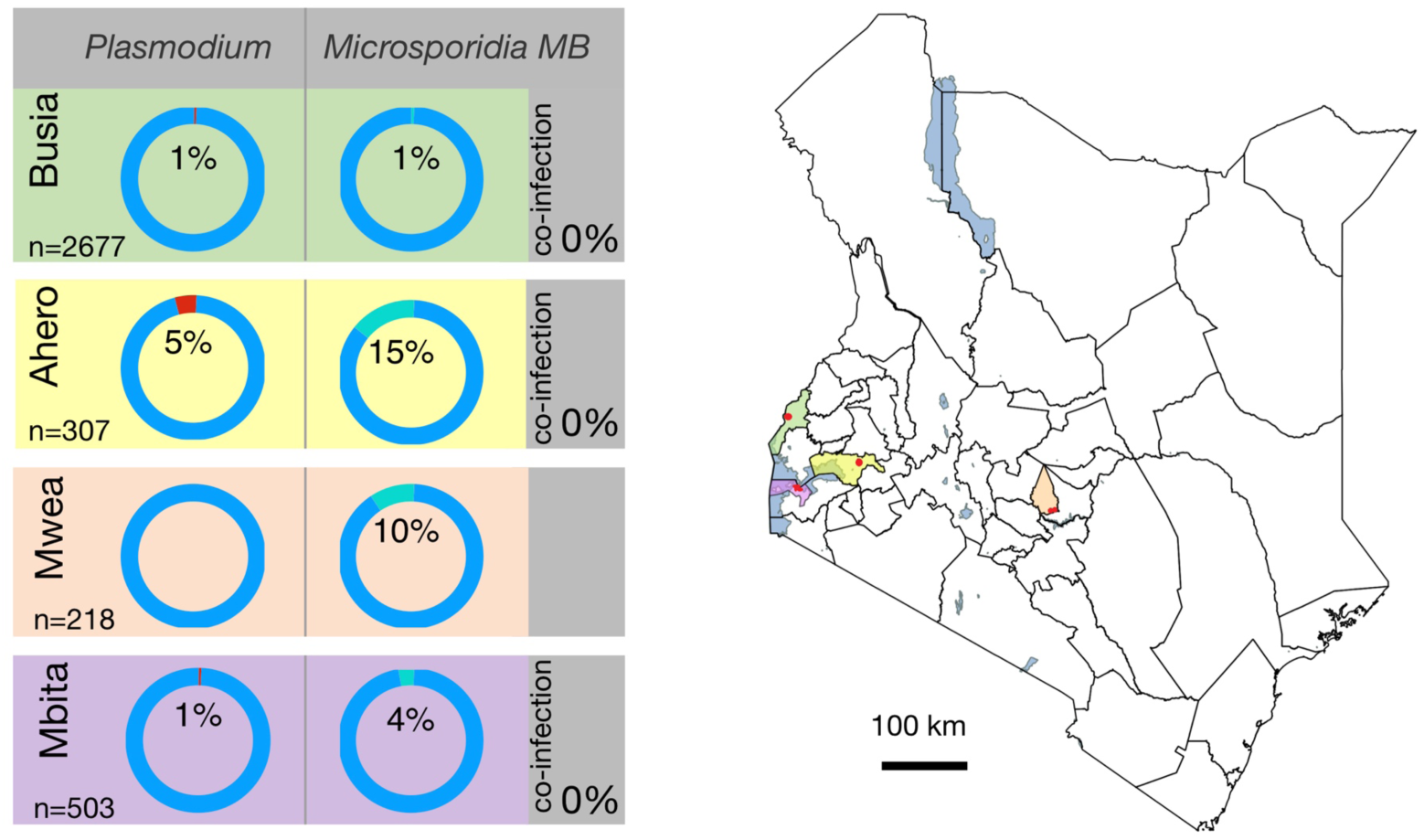
*Microsporidia MB* is found in geographically dispersed *An. arabiensis* populations. *Microsporidia MB* is found a prevalence range of 1-15% in *An. arabiensis* populations that also have a *Plasmodium* sporozoite infection prevalence range of 1-5%. *Microsporidia MB* and *Plasmodium* sporozoite co-infections were not observed. The color-coded locations of sampling sites are indicated on the map of Kenya.

**Extended Data Figure 3:**
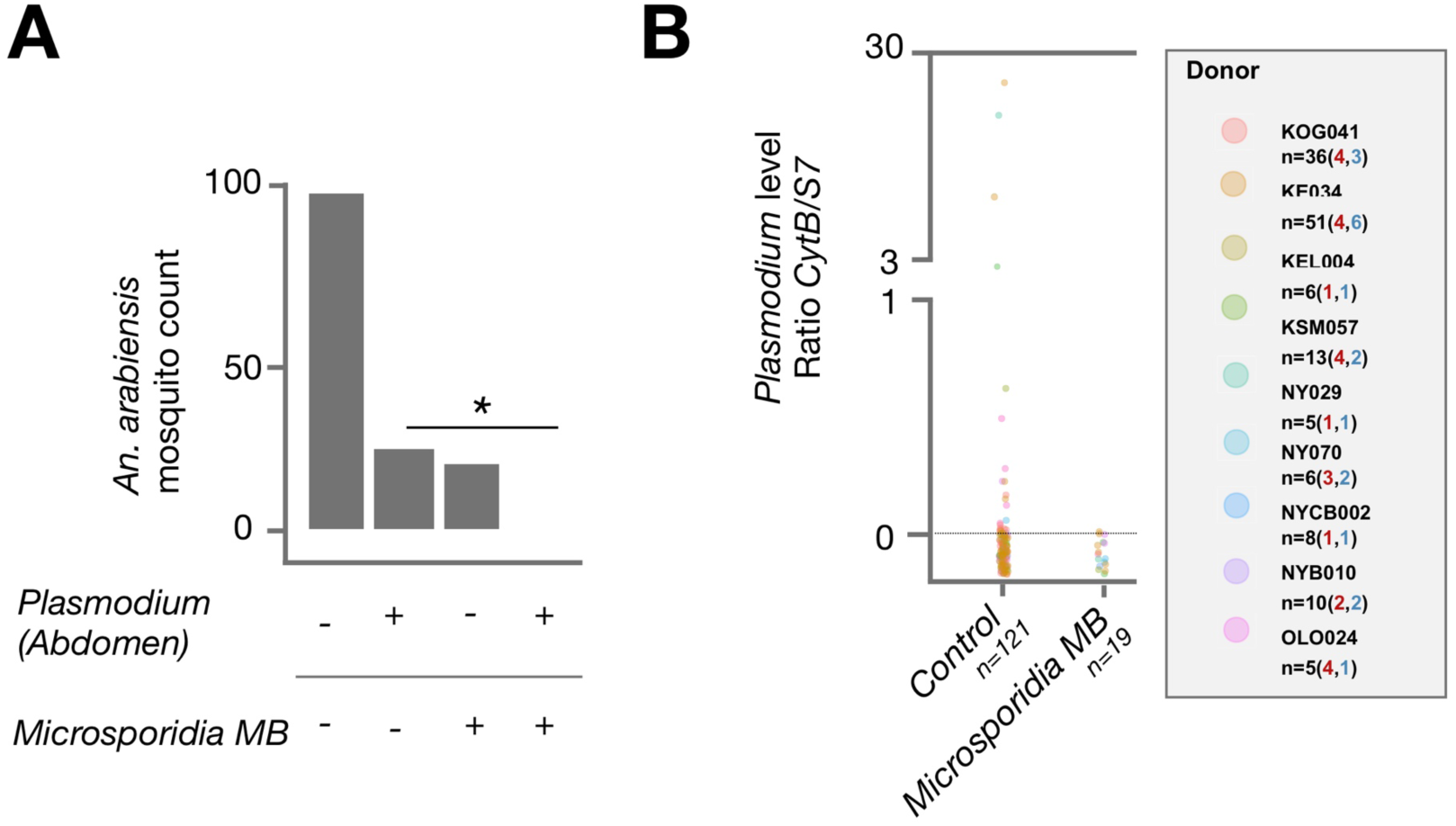
*Microsporidia MB* blocks oocyst formation in *An. arabiensis* after membrane feeding assay challenge with *P. falciparum.* The abdomen *Plasmodium* infection rate, reflecting presence of oocysts, in *Microsporidia MB* positive and *Microsporidia MB* negative mosquitoes. There was a significant absence of co-infected mosquitoes (two tailed fisher exact test, P=0.04 N=140).

**Extended Data Figure 4:**
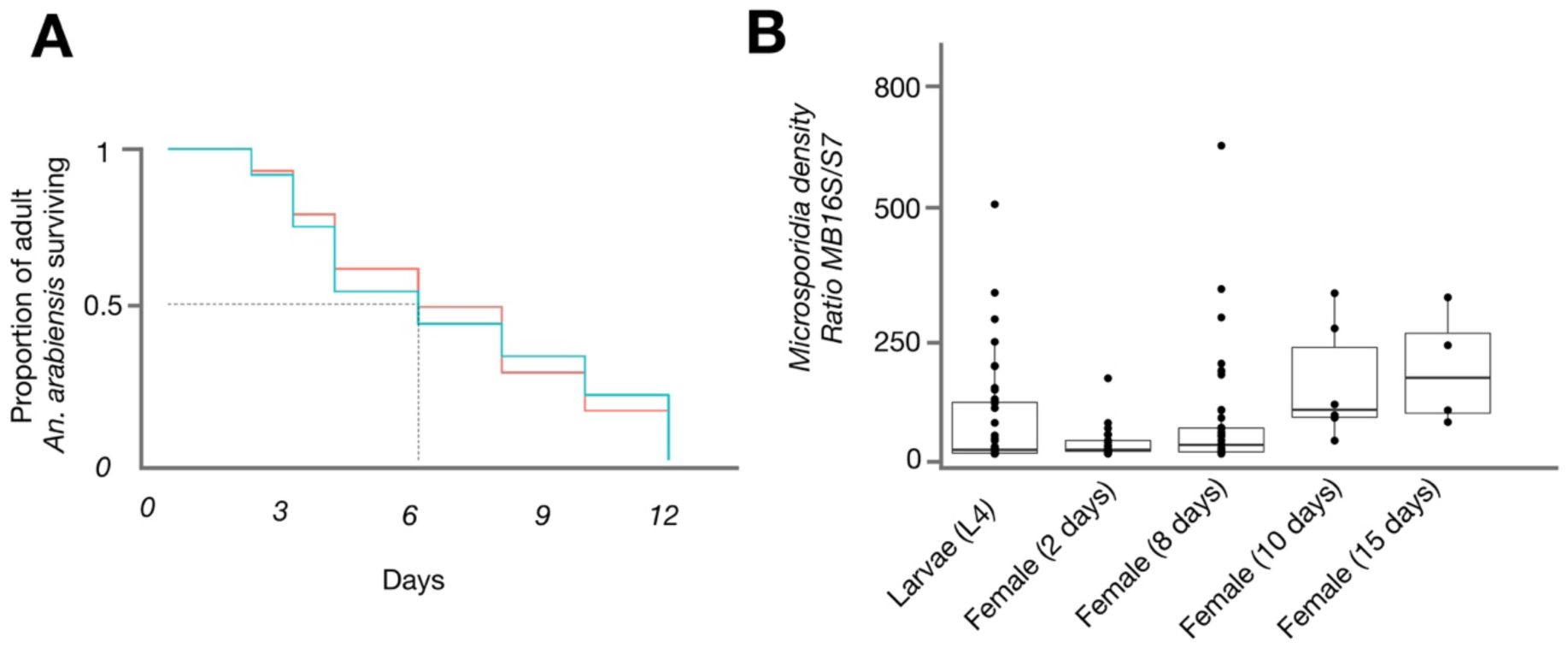
(A) The survival of adult G_1_ progeny of *Microsporidia MB* infected wild-caught females is not significantly different from uninfected counterparts, one experiment (ii) shown of three independent experiments (i-iii); (i, P=0.86, N>9, ii, P=0.97, N>41, ii, P=0.06 N>50, N denotes the minimum number of mosquitoes per condition). (B) The density of *Microsporidia MB* across different developmental stages. The density of *Microsporidia MB* is highly variable in larvae, lower in young adults but increase as adult *An. arabiensis* females age. Boxplot boundaries reflect the inter-quartile range. Each datapoint represents an individual mosquito from a total of five (L4-day 8) and three (day 10 and 15) independent experiments.

## Acknowledgments

We acknowledge Milcah Gitau of *icipe* Arthropod Rearing and Containment Unit for mosquito rearing assistance. We thank Ibrahim Kiche, Faith Kyengo and Ulrike Fillinger for assistance and advice.

## Author contributions

J.K.H conceived and designed the majority of the experiments. E.M and L.M. performed the majority of experiments. J.O and E.M. collected mosquitoes, screened them and prepared them for experiments. H.B. and M.V.M. carried out transmission-blocking and molecular identification experiments, respectively. J.K.H., S.P.S., E.M., L.M., M.V.M. and V.A.M. analyzed the data. J.K.H and S.P.S. wrote the manuscript.

